# Substrate rigidity suppresses the NRF2 antioxidant program in breast cancer cells

**DOI:** 10.64898/2026.11.23.768081

**Authors:** Safir Ullah Khan, Muhammad Danish Javed, Neha Roy Karintholil, Alfonso Méndez Tenorio, Dwayne G Stupack

**Affiliations:** Department of Obstetrics, Gynecology & Reproductive Sciences and Moores Cancer Center, University of California San Diego, La Jolla, CA, USA; Department of Material Sciences, University of California San Diego, La Jolla, CA, USA; Department of Biochemistry, Laboratory of Bioinformatics and Biotechnology, National Polytechnic Institute, Mexico City, Mexico

## Abstract

Matrix stiffness rises as breast tumors progress, but whether rigidity raises or lowers activity of the KEAP1-NRF2 antioxidant pathway is contested: work in breast cancer cells places the antioxidant response on soft matrix, while work in liver cancer places response on stiff. We used publicly available transcriptome data to ask the question across laboratories rather than within one system. Of eleven series identified, four dataset arms passed quality criteria fixed before any biological analysis. Across five contrasts spanning three cell lines and two matrix formats, one program responded in the same direction every time: the NRF2 antioxidant battery was lower on the more rigid substrate in all five. In contrast, SLC7A11, the cystine transporter supporting glutathione synthesis, decreased in all five, by up to 6.5-fold. YAP and TEAD targets did not track rigidity, rising on hydrogels and glass but falling on cell-derived matrix. NFE2L2 and its regulators were unchanged, while targets fell in proportion to their dependence on the antioxidant response element, pointing to control after transcription. An unsupervised partition made without reference to any pathway recovered the same program and survived removal of proliferation genes. In 1,097 TCGA and 1,980 METABRIC breast tumors, the matrix proxy was unrelated to the full battery until immune and hypoxic composition were accounted for, after which the same negative direction appeared in both; and a tumor-intrinsic score excluding macrophage-expressed members showed it more directly. Single-cell and spatial tests produced the predicted direction but could not resolve it, due to limited sample size and because the genes carrying the effect are largely undetectable at spot depth. Reduced antioxidant capacity on rigid substrate predicts greater sensitivity to agents killing through lipid peroxidation, a prediction we set out for experimental test.

**Author summary:** Tumors are stiffer than the tissue around them, and cancer cells sense that difference. Whether stiffness turns the cell’s main antioxidant defense up or down has been answered both ways, with studies in breast and in liver tissue reaching opposite conclusions. We reanalyzed gene expression data that other laboratories deposited in public repositories, because a direction that holds across many experiments is more informative than one measured in a single system.

The collected data require inspection before analysis. We judged each dataset against criteria set before looking at any biology and discarded three. In one excluded dataset, samples grown on soft gel detected roughly half as many genes as the other samples in the same experiment, a technical failure that would have been mistaken for a large biological effect.

In the datasets that were included, cells on stiffer surfaces consistently made less of the antioxidant machinery. The transcription factor controlling those genes was not itself reduced; only its targets were, which indicates that it is being held inactive rather than produced in smaller amounts. For breast cancer, this settles the direction, and it predicts that stiff tumor regions are more vulnerable to drugs that kill through oxidative damage.

## Introduction

The extracellular matrix of a breast tumor is stiffer than the tissue it replaces. Elastic moduli rise from a fraction of a kilopascal in normal mammary stroma to several kilopascals in invasive carcinoma, driven by collagen deposition and lysyl oxidase-mediated crosslinking [1,2]. Cells read this change. Substrate rigidity on its own is sufficient to redirect lineage commitment [3], and clinical elastography uses the same property to separate malignant from benign lesions [4].

How rigidity is converted into transcription is understood mainly through one axis. Nuclear translocation of YAP and TAZ scales with substrate stiffness and cytoskeletal tension, and their TEAD-dependent targets have become the standard readout of a mechanical response [5,6]. Matrix viscoelasticity, ligand presentation, and mechanosensitive ion channels all modulate the same machinery [7,8]. Since YAP/TAZ activity is an excellent marker, programs that respond to rigidity outside that axis are easily overlooked.

One such program controls redox balance. KEAP1 binds NFE2L2 (NRF2) in the cytoplasm and, as the substrate adaptor of a CUL3 ubiquitin ligase, directs it to the proteasome [9,10]. Oxidation of reactive KEAP1 cysteines halts that turnover; NRF2 accumulates and drives transcription from antioxidant response elements in genes for glutathione synthesis, NADPH supply, and thiol repair [11]. In tumors, the pathway follows both directions: it limits oxidative damage during transformation, and it also protects established cancer cells from chemotherapy and oxidative cell death [12–14].

Whether matrix rigidity raises or lowers NRF2 activity is unsettled, and the published answers point in opposite directions. In breast cancer cells, Romani and colleagues found that a soft matrix promotes DRP1-dependent mitochondrial fission, raises mitochondrial reactive oxygen species, activates NRF2, and thereby protects dormant metastatic cells from cisplatin [15]; soft substrate likewise raises reactive oxygen species and triggers protective autophagy in MDA-MB-231 cells [16]. In hepatocellular carcinoma, Elblova and colleagues reported the opposite on stiffness-graded collagen: cells held at 28 kPa showed lower reactive oxygen species, nuclear translocation of NRF2, and greater resistance to cisplatin and sorafenib than cells at 1.5 kPa [17]. Ferroptosis literature downstream of this axis splits in a similar manner. Stiffness has been reported to open a ferroptotic vulnerability through YAP-SKP2 signaling [18] and, in lung adenocarcinoma, to close one through an integrin-Hippo-ABCG1 axis [19]; stiffness-dependent control of xCT has been described in liver cancer [20], stiffness-induced redox signaling in pancreatic cancer [21], and mechanically induced autophagy as a ferroptosis sensitizer [22].

The disagreement has consequences. NRF2 output sets the ceiling on cystine import through SLC7A11 and on the glutathione available to GPX4, the two constraints that decide whether a cell survives lipid peroxidation [23–26], and drug-tolerant persister states depend on the same axis [27]. If rigidity lowers NRF2 output, stiff tumor regions should be more sensitive to ferroptosis inducers; if it raises NRF2 output, they should be less sensitive. The studies above differ in tissue of origin, in the stiffness range tested, in matrix format, and in readout, since nuclear localization of a protein and steady-state transcript levels of its targets are not the same measurement. No single experiment separates these explanations.

Transcriptome data from stiffness experiments have accumulated in public repositories [28], which makes it possible to ask whether a direction holds across laboratories, cell lines and matrix formats rather than within one system. Reuse of these data is not automatic. Deposited series differ in design, and some carry technical failures large enough to produce a confident result that is entirely artifactual. We therefore screened every candidate series against quality criteria fixed before any biological analysis, analyzed only what survived, and asked which programs respond to rigidity reproducibly. One did.

## Results

### Most publicly available stiffness datasets do not meet basic quality criteria

We queried GEO for transcriptome studies comparing breast cancer cells cultured on substrates of differing stiffness and catalogued eleven series. These hold 137 deposited samples, of which 122 are human; the remaining 15 are the mouse MMTV-PyVT arm of GSE107063, deposited on a separate platform and excluded at screening because the first criterion requires a human breast cancer line. Six series, contributing seven dataset arms, had the design and data type required for a direct comparison. Each arm was evaluated against two criteria applied before any biological analysis (Fig 1A and 1B): replicates within a condition were required to correlate at Spearman ρ ≥ 0.9, and the median within-group correlation was required to exceed the median between-group correlation by at least 0.02.

**Fig 1.**
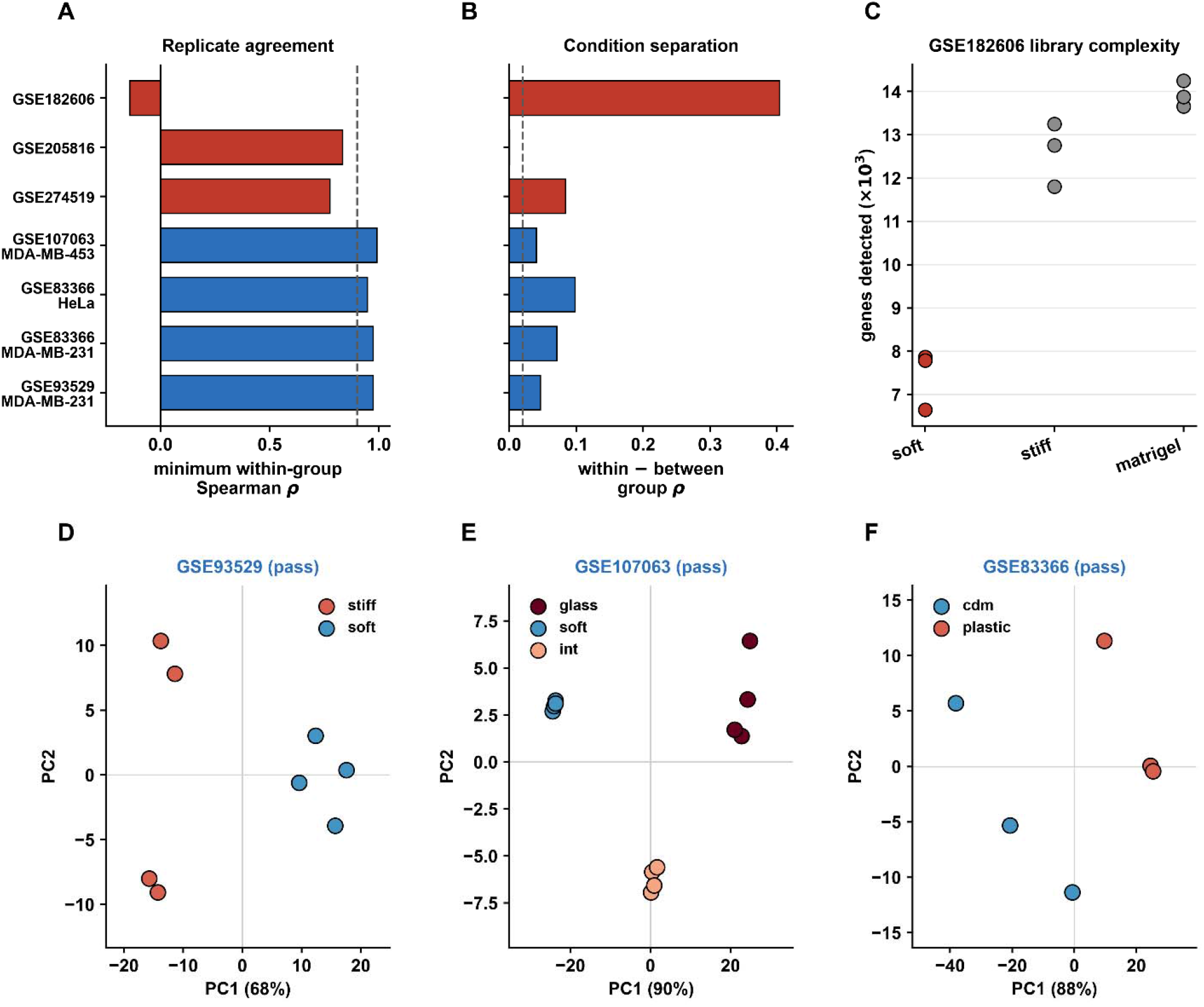
Assembly of the compendium and quality control. (A) Minimum within-group Spearman correlation across all replicate pairs, for the seven dataset arms whose design permitted a direct comparison; the dashed line marks the retention threshold of 0.90. (B) Condition separation, defined as the median within-group correlation minus the median between-group correlation; the dashed line marks the threshold of 0.02. Series failing either criterion are shown in red, retained series in blue. (C) Genes detected per sample in GSE182606 at matched sequencing depth: 6,644 to 7,865 on compliant substrate, 11,806 to 13,248 on stiff substrate and 13,655 to 14,247 on Matrigel. (D-F) Principal component analysis of three of the four retained arms: GSE93529 (n = 8), GSE107063 (n = 12) and the MDA-MB-231 arm of GSE83366 (n = 6); the HeLa arm of GSE83366 behaves equivalently and is tabulated in S1 Table. Substrate conditions separate along PC1 in every case, accounting for 68%, 90%, and 88% of total variance respectively.

Four datasets met both criteria: GSE93529 [29] (MDA-MB-231, 0.7 kPa hydrogel versus plastic, n = 8, minimum within-group ρ = 0.97), GSE107063 [30] (MDA-MB-453, 5 kPa, 30 kPa and glass, n = 12, ρ = 0.99), and the MDA-MB-231 and HeLa arms of GSE83366 [31] (cell-derived matrix versus plastic, n = 6 each, ρ = 0.97 and 0.95).

Three failed, each for a distinct reason. In GSE182606 [32] the compliant-substrate replicates were anticorrelated (ρ = −0.14) and detected 6,644–7,865 genes, against 11,806–13,248 on stiff substrate and 13,655–14,247 on Matrigel at matched depth (Fig 1C); this is a library complexity failure that would be indistinguishable from a large biological effect in downstream analysis. GSE205816 [33] showed negligible separation between conditions (within minus between ρ = 0.0008). GSE274519 fell below the replicate agreement threshold (ρ = 0.78). Five further series were excluded on design grounds, including one deposited solely as an unannotated de novo assembly in which none of 76,445 loci were shared across all nine samples. All inclusion and exclusion decisions, with reasons, are recorded sample by sample in S1 Table. Principal component analysis separated conditions on PC1 in each retained dataset; three are shown (Fig 1D–1F).

### The stiffness response is predominantly cell-line specific

Differential expression was computed for five contrasts (Fig 2E). Genes reaching FDR < 0.05 ranged from 1,315 (HeLa, cell-derived matrix) to 6,213 (MDA-MB-453, glass). Median absolute effect sizes were modest on hydrogels (0.24–0.33 log2 units) and larger on cell-derived matrix (0.65–0.77) (Fig 2F). Throughout, a positive log2 fold change denotes higher expression on th stiffer or more rigid substrate.

**Fig 2.**
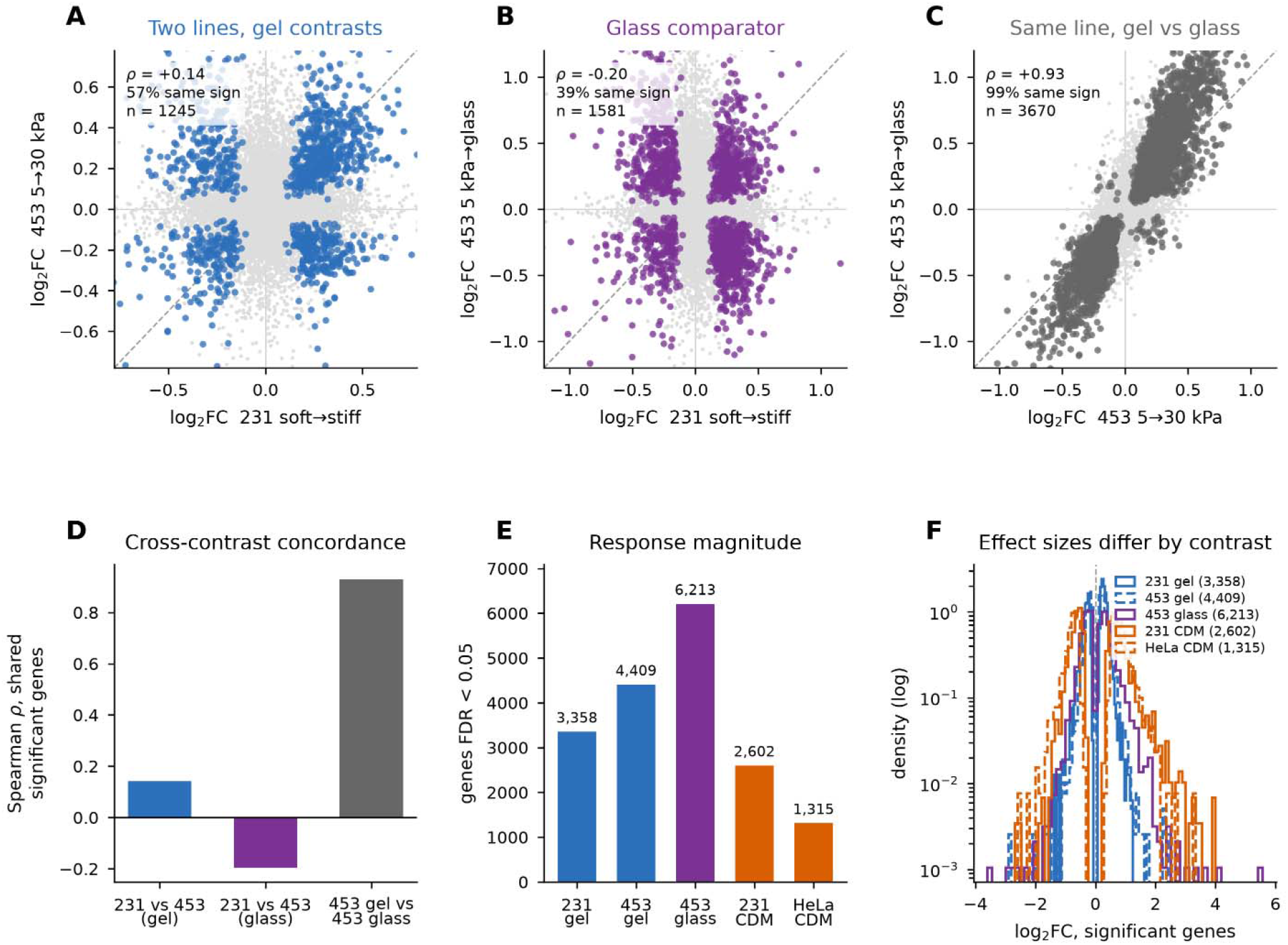
The transcriptional response to rigidity is largely cell-line specific. (A-C) Log2 fold changes compared between pairs of contrasts. Genes significant at FDR < 0.05 in both contrasts are highlighted; all other measured genes are shown in gray. Spearman ρ, the fraction of shared genes changing in the same direction, and the number of shared genes are shown in the inset. (A) MDA-MB-231 hydrogel against MDA-MB-453 5 to 30 kPa (ρ = +0.14, 57.2% same sign, n = 1,245). (B) MDA-MB-231 hydrogel against MDA-MB-453 5 kPa to glass (ρ = −0.20, 39.1%, n = 1,581). (C) The two MDA-MB-453 contrasts against one another (ρ = +0.93, 98.8%, n = 3,670); these share a cell line and differ only in the rigid comparator and so serve as a positive control. (D) The three correlation coefficients side by side. (E) Genes reaching FDR < 0.05 in each contrast. (F) Distribution of log2 fold change among significant genes, by contrast. Throughout, a positive log2 fold change denotes higher expression on the stiffer or more rigid substrate.

Agreement between cell lines was weak. Among 1,245 genes significant in both the MDA-MB-231 and MDA-MB-453 hydrogel contrasts, Spearman ρ was +0.14 with 57% directional agreement (Fig 2A). Against the MDA-MB-453 glass contrast, the relationship inverted (ρ = −0.20, 39% agreement, n = 1,581; Fig 2B). Both correlations reached significance only by virtue of sample size (P < 1 × 10□³□□).

The two MDA-MB-453 contrasts agreed closely with one another (ρ = +0.93, 98.8% directional agreement, n = 3,670; Fig 2C). As these share a cell line and differ only in the rigid condition used, this serves as a positive control: the weak cross-line agreement reflects genuine cell-line specificity rather than measurement noise (Fig 2D).

### A single module is suppressed by rigidity in every contrast

We next asked whether any coherent program behaved consistently despite weak gene-level agreement. The NRF2 antioxidant battery was reduced in all five contrasts, with mean log2 fold changes of −0.25, −0.68, −1.04, −0.44 and −0.23 (Fig 3A). Seven of the fourteen battery genes were negative in all five contrasts, including SLC7A11 (−0.21, −2.07, −2.70, −1.40, −1.01), GCLM, NQO1, TXNRD1, SRXN1, HMOX1, and ME1; an eighth, AKR1C1, was negative in each of the three contrasts in which it was measured.

**Fig 3.**
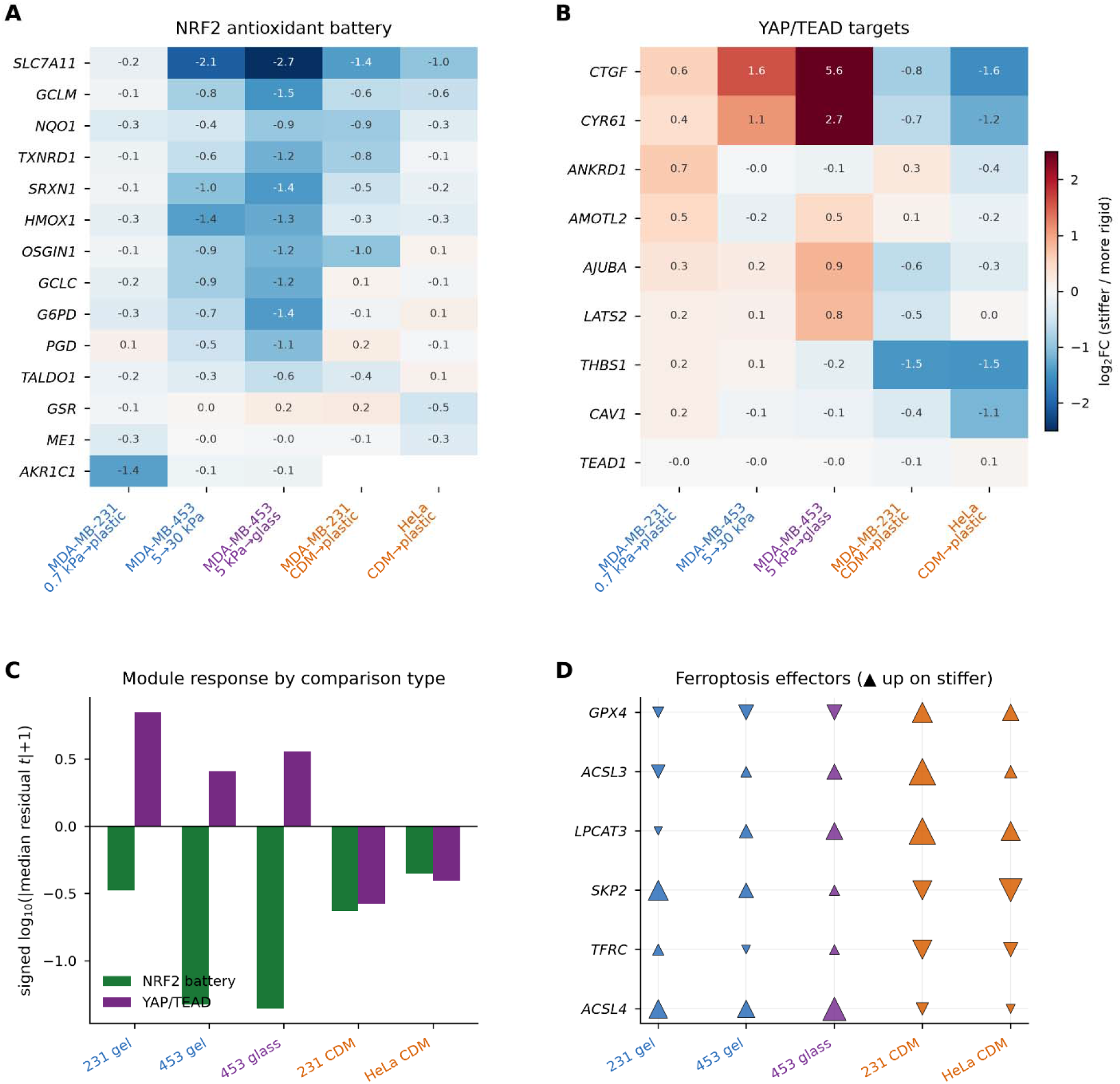
The NRF2 battery responds in one direction in every contrast; YAP/TEAD targets do not. (A) Log2 fold changes of the 14 NRF2 battery genes across the five contrasts, with column means of −0.25, −0.68, −1.04, −0.44 and −0.23. Seven genes are negative in all five contrasts; AKR1C1, which the two cell-derived-matrix arrays do not measure, is negative in each of the three contrasts in which it is measured. (B) The same display for nine YAP/TEAD target genes; column means +0.33, +0.31, +1.11, −0.46 and −0.69, with CTGF ranging from +5.57 on glass to −1.56 in HeLa on cell-derived matrix. (C) Module response per contrast, given as the signed log□□ of the median residualized t statistic for module members, for the NRF2 battery and YAP/TEAD targets. The NRF2 battery is negative in 5 of 5 contrasts and YAP/TEAD targets in 2 of 5. (D) Log2 fold changes of six ferroptosis effectors, shown as upward triangles where expression is higher on the stiffer substrate and downward triangles where it is lower, with marker area scaled to effect size; GPX4 tracks with ACSL4 rather than opposing it. Positive values denote higher expression on the stiffer or more rigid substrate.

YAP/TEAD targets did not behave in this pattern (Fig 3B). They increased on hydrogel and glass (means +0.33, +0.31, +1.11) and decreased on cell-derived matrix (−0.46, −0.69), with CTGF ranging from +5.57 on glass to −1.56 in HeLa on cell-derived matrix. Testing each module against all remaining measured genes confirmed the dissociation: NRF2 was negative in 5 of 5 contrasts, YAP/TEAD in 2 of 5, and the ferroptosis arm in 3 of 5 (Fig 3C). Individual ferroptosis effectors showed no consistent direction; notably GPX4, which protects against ferroptosis, tracked with ACSL4 rather than opposing it (Fig 3D).

### Suppression scales with promoter dependence and spares the pathway itself

Reduced NRF2 production would predict lower NFE2L2 transcript levels. This was not observed. Averaged across contrasts, NFE2L2, KEAP1, CUL3 and MAFG changed by −0.02 log2 units (n = 18 values, SD 0.12), whereas six strongly ARE-dependent targets decreased by −0.78 (n = 30, SD 0.63; Mann-Whitney P = 1.5 × 10□□) (Fig 4B).

**Fig 4.**
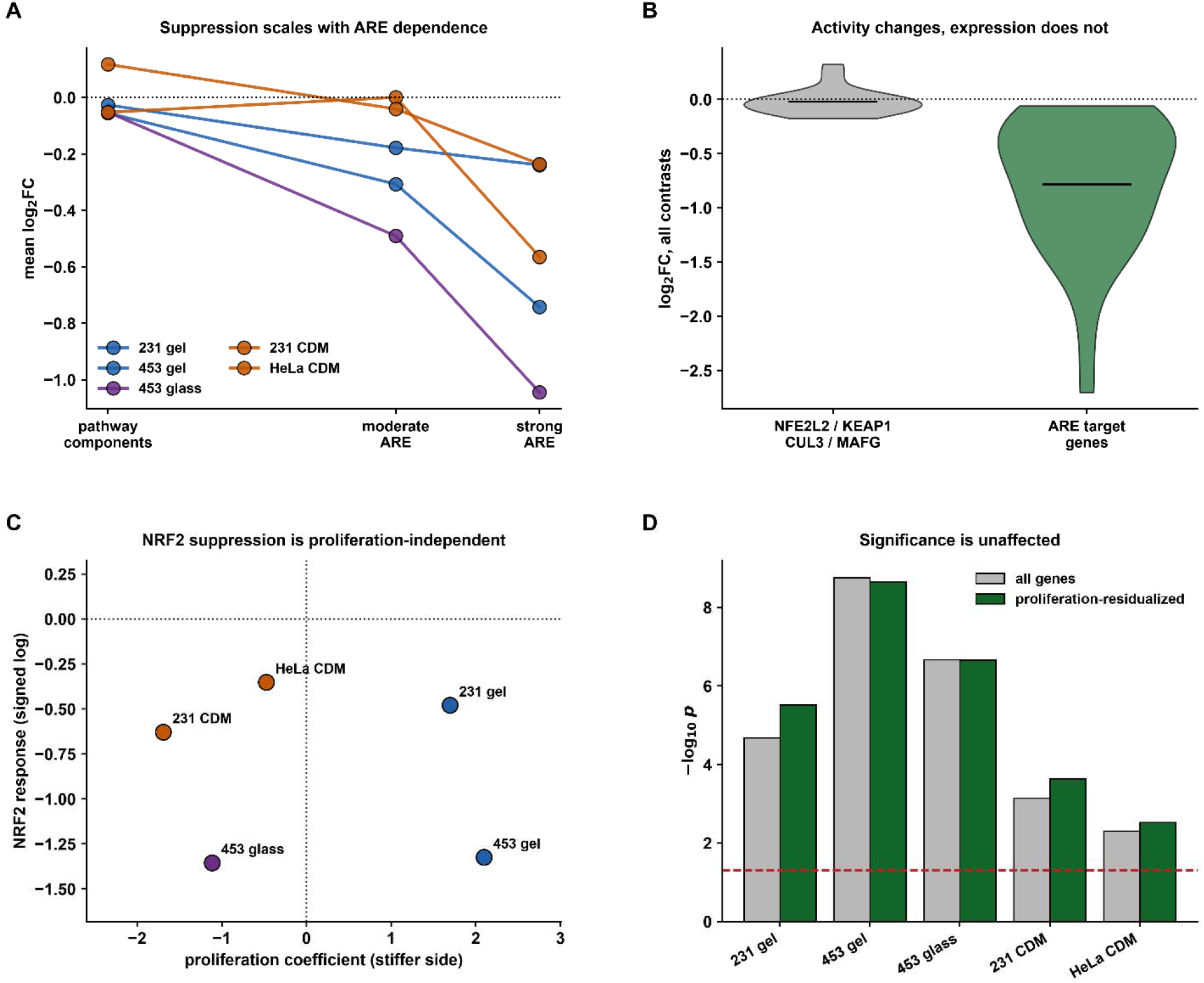
Suppression scales with promoter dependence and does not follow proliferation. **(A)** Mean log2 fold change of NRF2 battery genes by antioxidant response element dependence, assigned before analysis to three tiers: pathway components (NFE2L2, KEAP1, CUL3, MAFG; 4 genes), moderately dependent targets (8 genes), and strongly dependent targets (11 genes). The gradient is present in four of five contrasts, all but MDA-MB-231 on cell-derived matrix. (B) Pooled log2 fold changes of the four pathway components (n = 18 values, mean −0.02) against the six canonical antioxidant response element targets SLC7A11, NQO1, TXNRD1, SRXN1, GCLM and HMOX1 (n = 30, mean −0.78); two-sided Mann-Whitney U = 517, P = 1.5 × 10□□. Pooling all eleven strongly dependent genes instead gives n = 53 and a mean of −0.57. (C) NRF2 module response against the proliferation coefficient of the ordinary least squares model, one point per contrast; the proliferation coefficient (+1.70, +2.10, −1.11, −1.69, −0.47) changes sign across contrasts, while the NRF2 response does not. (D) Significance of the NRF2 module before and after residualization on proliferation; every contrast remains significant, with a largest P of 3.0 × 10□³.

Stratifying NRF2 pathway genes by antioxidant response element dependence produced a graded effect (Fig 4A). This tiering covers 23 genes, the fourteen battery members together with the four pathway components and five further targets, and therefore is wider than the battery in Fig 3A. Pathway components changed by −0.05 to +0.12 across contrasts, moderately dependent genes by 0.00 to −0.49, and strongly dependent genes by −0.24 to −1.05. The gradient was present in four of five contrasts.

Because proliferation rate varies with substrate, we tested whether it accounted for the result. The proliferation coefficient changed sign across contrasts (+1.70, +2.10, −1.11, −1.69, −0.47) while the NRF2 response remained negative in all five (Fig 4C). After removing proliferation genes and retesting, every contrast remained significant (largest P = 3.0 × 10□³) (Fig 4D). A covariate that reverses direction while the effect does not cannot be driving that effect.

### An unsupervised partition recovers the same program

The preceding analysis begins from a predefined gene set, which raises the possibility that the set was selected with the result in view. We therefore repartitioned the data without reference to any pathway.

Genes significant at FDR < 0.05 in both contrasts of a comparison were assigned to one of four quadrants by the sign of their response in MDA-MB-231 and MDA-MB-453 (Fig 5A), and the hydrogel and glass arms of the MDA-MB-453 comparison were intersected within each quadrant (Fig 5B–5F). Ribosomal, mitochondrial, and small RNA symbols were removed beforehand (159 symbols), leaving 16,953 genes measured in all three contrasts as the background. The two MDA-MB-453 arms agreed substantially and uniformly, sharing 158 to 300 transcripts per quadrant (Jaccard 0.45–0.59; all P < 1 × 10□²□³).

**Fig 5.**
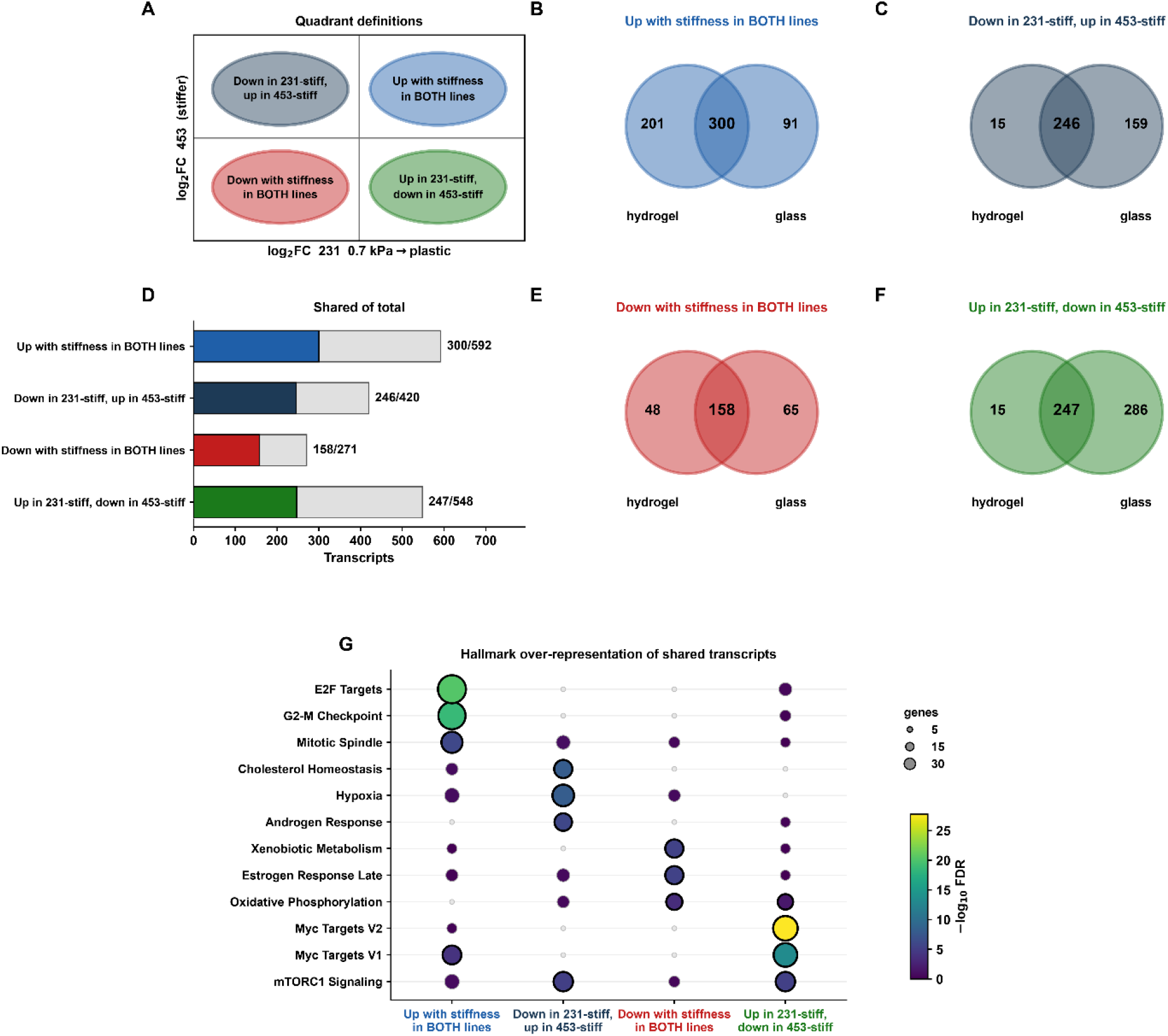
A partition made without reference to any pathway recovers the same program. **(A)** Schematic of the quadrant definitions: genes significant at FDR < 0.05 in both contrasts of a comparison are assigned to four quadrants by the sign of their response in MDA-MB-231 and in MDA-MB-453. (B, C, E, F) Venn diagrams of the overlap between the hydrogel and glass arms of the MDA-MB-453 comparison within each quadrant, against a background of 16,953 genes measured in all three contrasts. Shared transcripts and Jaccard indices: Q1 300 and 0.51, Q2 246 and 0.59, Q3 158 and 0.58, Q4 247 and 0.45; every overlap reached P < 1 × 10□²□³ by one-sided hypergeometric test. (D) Shared transcripts as a fraction of the union of the two arms, by quadrant: 300/592, 246/420, 158/271 and 247/548. (G) Hallmark terms reaching FDR < 0.05 in each quadrant by over-representation analysis. Q3, suppressed by rigidity in both cell lines, carries xenobiotic metabolism, oxidative phosphorylation, fatty acid metabolism, glycolysis, and the reactive oxygen species pathway. Q1, induced in both lines, carries cell-cycle terms almost exclusively.

Each quadrant carried a distinct program (Fig 5G, S1 Fig). Transcripts suppressed by rigidity in both cell lines were enriched for xenobiotic metabolism (12 of 158 genes, 6.7-fold, FDR = 3.8 × 10□□), oxidative phosphorylation (5.7-fold), fatty acid metabolism, glycolysis and the reactive oxygen species pathway (9.5-fold, FDR = 3.9 × 10□³). This recovers the NRF2 battery without it having been specified. Transcripts induced in both lines were almost exclusively cell cycle (E2F targets 9.7-fold, FDR = 5.4 × 10□²¹; G2-M checkpoint 9.3-fold). The two discordant quadrants carried cholesterol homeostasis and hypoxia programs (12.0-fold and 6.5-fold, both FDR = 7.6 × 10□□), and ribosome biogenesis with MYC targets (MYC targets V2 29.4-fold, FDR = 1.8 × 10□²□), respectively.

Repeating the analysis with 671 proliferation genes removed from both the query set and the background separated these outcomes cleanly (S2 Fig). The suppressed-in-both quadrant contained 3 proliferation genes of 158 (1.9%), and all eight leading terms held or strengthened. The induced-in-both quadrants contained 59 of 300 (19.7%), and its cell-cycle terms fell below significance, leaving only an mRNA splicing program (FDR = 1.1 × 10□□). Across 32 term comparisons, 28 held, 3 weakened, and 1 was lost.

### In tumor tissue the association is masked by immune and hypoxic composition

We next asked whether the same direction is detectable in human tumors. A score measured in bulk tissue mixes tumor-intrinsic signal with infiltrate, because several battery members, HMOX1 in particular, are expressed by tumor-associated macrophages. We therefore specified three NRF2 scores in advance of testing: the full fourteen-gene battery, the ten antioxidant response element-dependent members, and a seven-gene tumor-intrinsic score comprising the strongly dependent members of Fig 4A with HMOX1 removed as myeloid-expressed and AKR1C1 removed as unreliably detected (SLC7A11, GCLM, NQO1, TXNRD1, SRXN1, OSGIN1, GCLC). The seven-gene score is the primary outcome throughout this section; all three are reported for every test in S1 Table.

In 1,097 TCGA-BRCA primary tumors [34], a matrix stiffness proxy built from collagen and lysyl oxidase transcripts was at most weakly related to the primary score (Spearman ρ = −0.068, P = 0.025; Fig 6A), and unrelated to the full battery (ρ = +0.014, P = 0.65). The proxy was not independent of tissue composition. It correlated with myeloid content (ρ = +0.36, P = 3.5 × 10□³□) and with a hypoxia score built from ten canonical HIF1A targets [35] (ρ = +0.15, P = 1.1 × 10□□), both of which raise an NRF2 score, while epithelial content was unrelated to the proxy (ρ = −0.004, P = 0.90) (S3 Fig). Positively correlated inputs of this kind suppress any underlying negative association, completely for the full battery and partially for the tumor-intrinsic score.

**Fig 6.**
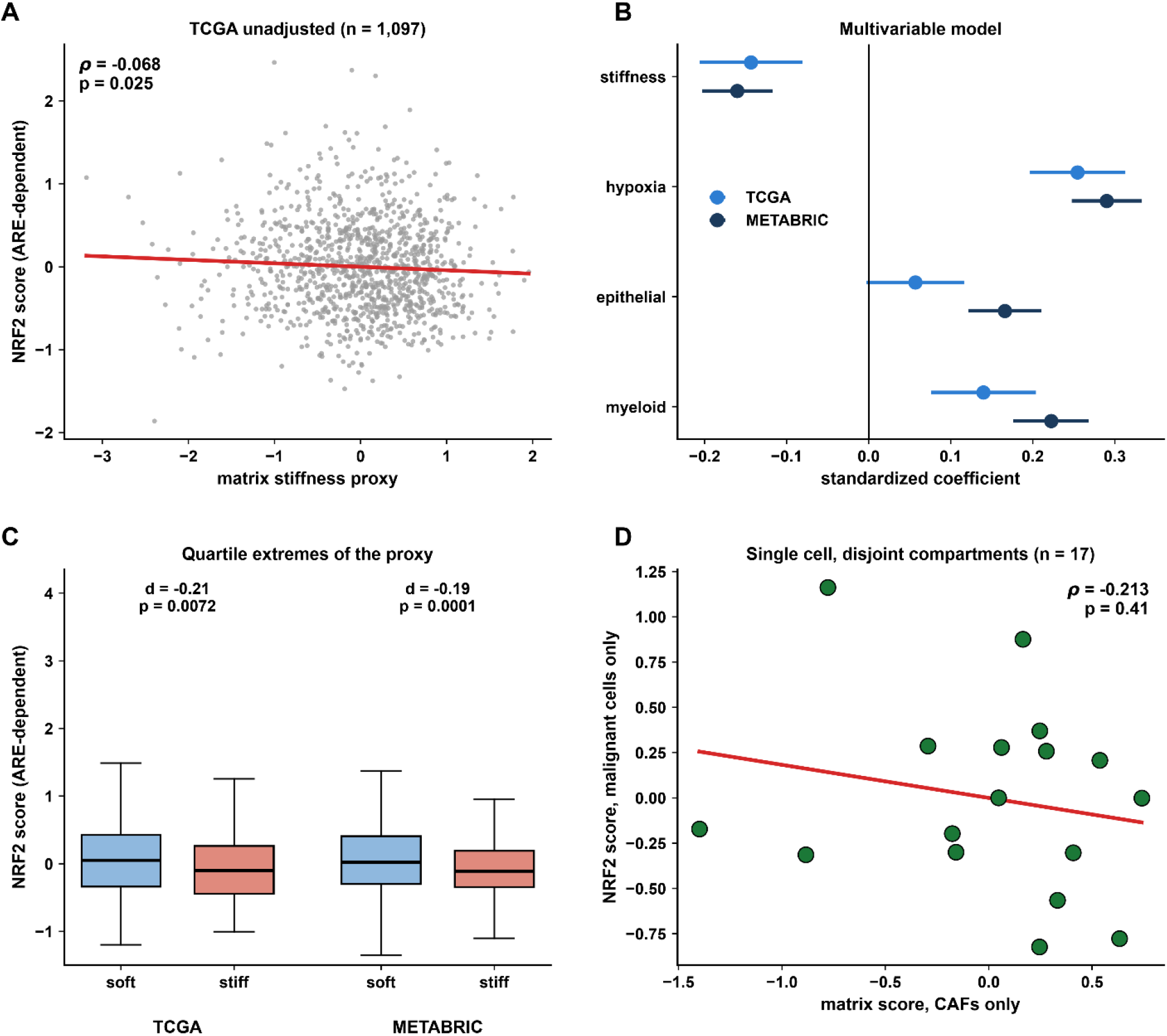
The direction is recovered in human tumors once tissue composition is accounted for. (A) Matrix stiffness proxy against the seven-gene tumor-intrinsic NRF2 score across 1,097 TCGA-BRCA primary tumors; the marginal relationship is weak (Spearman ρ = −0.068, P = 0.025). (B) Standardized coefficients with 95% confidence intervals from the model regressing that score on the stiffness proxy, hypoxia, epithelial content and myeloid content, in TCGA (n = 1,097) and METABRIC (n = 1,980). The stiffness term is negative in both cohorts (β = −0.143 and −0.160) while all three covariates are positive. (C) The same score in the top and bottom quartiles of the proxy, by cohort (TCGA d = −0.21, P = 7.2 × 10□³; METABRIC d = −0.19, P = 1.0 × 10□□). (D) Single-cell test, with the NRF2 score computed in malignant cells only against the matrix score computed in cancer-associated fibroblasts only, one point per tumor, across the 17 tumors contributing at least 20 cells to both compartments (ρ = −0.213, P = 0.41).

In a model including hypoxia, epithelial and myeloid content, the stiffness proxy was a significant negative predictor of the tumor-intrinsic score (β = −0.143 per standard deviation, P = 8.2 × 10□□), with hypoxia at +0.25, myeloid at +0.14 and epithelial at +0.06 (P = 0.061) (Fig 6B). A rank-based partial correlation gave ρ = −0.147 (P = 1.0 × 10□□), and tumors in the top and bottom quartiles of the proxy differed by d = −0.21 (P = 7.2 × 10□³). Specifying this model in advance, we repeated it in 1,980 METABRIC tumors [36,37]. The estimate replicated at β = −0.160 (P = 3.9 × 10□¹³), every covariate reproduced in sign, the partial correlation gave ρ = −0.186 (P = 7.5 × 10□¹□), and quartile extremes differed by d = −0.19 (P = 1.0 × 10□□) (Fig 6C). On the full fourteen-gene battery, the adjusted estimate is of the same size in both cohorts (β = −0.157 and −0.162), but the quartile contrast is null (d = −0.02 and −0.06), as expected if infiltrate-derived expression obscures a tumor-intrinsic effect at the margins of the proxy. The reduction in the myeloid coefficient from +0.35 on the full battery to +0.14 on the seven-gene score is itself evidence that the removed members were carrying infiltrate signal.

Two higher-resolution tests were attempted, and neither was adequately powered. In 26 breast tumors profiled by single-cell RNA sequencing [38], scoring the battery in malignant cells and matrix genes in cancer-associated fibroblasts, which are disjoint populations and so cannot be coupled by composition, gave ρ = −0.213 across the 17 tumors contributing at least 20 cells to both compartments (P = 0.41; Fig 6D). The direction follows a predicted trend, but power at this effect size and sample size is 0.13, therefore, the test discriminates minimally. In six matched spatial transcriptomic sections, the association was null once sequencing depth was controlled (β = +0.010, P = 0.72). The limitation is measurement rather than sample size: SLC7A11 was detected in only 0.5 to 8.7% of spots, so the antioxidant response element-dependent arm is largely unmeasurable at that depth. Consistent with this, the six battery genes lost to single-cell detection carry roughly twice the culture effect of the eight retained (mean −0.76 against −0.38, P = 0.024), which recovers the promoter-dependence gradient of Fig 4 by a partition that makes no reference to promoter architecture (S3 Fig).

## Discussion

Across five contrasts drawn from three independent studies, spanning three cell lines and two matrix formats, the NRF2 antioxidant battery was lower on the more rigid substrate every time. No other module tested behaved this way, and an unsupervised partition that began from no pathway at all returned the same genes. For breast cancer cells in culture, the direction of stiffness-redox coupling is therefore the one reported on soft matrix, not the one reported on stiff.

This agrees with the functional work. Romani and colleagues showed that soft matrix drives DRP1-dependent mitochondrial fission, raises mitochondrial reactive oxygen species and activates NRF2 in breast cancer cells, and that knocking down NRF2 restored cisplatin sensitivity in dormant lung metastases [15]. Soft substrate also raises reactive oxygen species in MDA-MB-231 cells, where protective autophagy limits the resulting apoptosis [16]. These studies measured protein localization, reactive oxygen species, and drug response in one system each; we measured steady-state transcript levels of the downstream battery in three systems that none of them used. The two kinds of evidence are independent, and they are directionally consistent. The agreement is the strongest claim this reanalysis can make, because the datasets were deposited by laboratories asking other questions.

Work in liver cancer points in an opposing direction [20], yet the difference is informative rather than simply contradictory. Elblova and colleagues found that hepatocellular carcinoma cells on 28 kPa collagen showed lower reactive oxygen species, nuclear NRF2, and resistance to cisplatin and sorafenib relative to 1.5 kPa, with KEAP1 falling at 4 h and rising above the soft condition by 48 h [17]. Four differences separate this experiment from ours. The tissue is different. The stiffness range is narrow and entirely within the physiological span, whereas four of our five contrasts use plastic or glass, which are three orders of magnitude stiffer than any tissue [39]. The readout is different: nuclear translocation of a protein at a single time point is not equivalent to steady-state transcript abundance of its targets, and a pathway under post-translational control can change in one without changing in the other. In addition, the KEAP1 response that Elblova and colleagues report is itself biphasic in time, so a transcriptional snapshot taken at one time point could land on either side of it. The ferroptosis literature splits along similar lines, with stiffness reported to open a ferroptotic vulnerability in one setting [18] and close one in another [19–22]. We would not generalize our result beyond breast cancer cells, and the liver data give a concrete reason not to.

The mechanism our data support is control after transcription. NFE2L2 and its regulators KEAP1, CUL3 and MAFG were flat, averaging −0.02 log2 units across 18 values, while targets fell in proportion to how strongly their promoters depend on the antioxidant response element. Lower production of the transcription factor would not produce that gradient; lower activity of a constant pool would. This is the behavior expected of the KEAP1-CUL3 degradation switch [9,10], and it fits the fission mechanism directly: if soft matrix raises mitochondrial reactive oxygen species and thereby releases NRF2 [15], then rigid substrate should lower the input to that switch, leaving pathway transcripts untouched and antioxidant response element output reduced. Our data are consistent with that account but cannot establish it, since we measured neither NRF2 protein nor reactive oxygen species.

The redox response also dissociated from the canonical mechanotransduction readout. YAP and TEAD targets rose on hydrogels and glass and fell on cell-derived matrix, with CTGF spanning +5.57 to −1.56 across contrasts, while the NRF2 battery fell in all five. Substrate rigidity is therefore not a single program read out by a single reporter, and a study that scores the mechanical response by YAP/TAZ activity alone [5,6] can be insensitive to a redox change moving in a fixed direction underneath it. Metabolic programs under mechanical control behave similarly [40].

In tumor tissue the direction is present but partly masked. Taken marginally, the stiffness proxy and the full fourteen-gene NRF2 score are unrelated in both cohorts, and the proxy covaries with myeloid content and with hypoxia, both of which raise the score. HMOX1 and other battery members are expressed by tumor-associated macrophages, so a bulk measurement mixes tumor-intrinsic signal with infiltrate. Restricting the score to seven members not subject to that contamination makes the association visible even marginally, and the myeloid coefficient falls by more than half, which is what should happen if the removed genes were carrying infiltrate signal. Once composition is accounted for, the direction observed in culture is clear in both cohorts, at β = −0.143 in TCGA and −0.160 in METABRIC, with the model fixed before the replication cohort was examined and every covariate reproducing in sign. This is in part a suppression effect and should be read with the caution that implies: the estimate is small, it exists relative to a covariate set, and a transcriptional stromal score indexes desmoplastic content rather than tissue modulus. The two higher-resolution designs were uninformative rather than contrary. Comparing disjoint single-cell compartments gave the predicted direction across 17 tumors but at a power of only 0.13, and the spatial assay cannot detect the antioxidant response element-dependent genes at spot depth, where SLC7A11 appears in fewer than 9% of spots. No human analysis pointed the other way. Settling the question in tissue requires stiffness measured directly, by elastography or atomic force microscopy, paired with expression from the same region [4].

Several limitations bound the claim. Three cell lines contribute, and one of them, HeLa, is not a breast line; it is retained because it provides a second, independent cell-derived matrix contrast, and it behaves like the others. Plastic and glass are not physiological comparators [39], and cell-derived matrix differs from plastic in ligand density and architecture as well as in stiffness [41], so the contrasts are not clean manipulations of modulus alone. The evidence is transcriptional throughout; no protein, flux or reactive oxygen species measurement is included. One series, GSE93529, has a modulus discrepancy between its repository record and its publication, noted in S1 Table. Two probe-collapsing rules were used in different parts of the analysis, each applied uniformly within its own analysis but differing between them, which we state in Methods rather than conceal. Three NRF2 scores were evaluated in the human cohorts, and the extreme-group testing was exploratory; the primary score was chosen on biological grounds, and all seventeen comparisons are tabulated. Finally, four dataset arms is a small compendium; it is small because most deposited series did not survive quality control, which is itself a result worth reporting.

The prediction follows directly. NRF2 output sets cystine import through SLC7A11 and the glutathione supply to GPX4, the limiting constraints on survival of lipid peroxidation [23–26]. SLC7A11 fell in all five contrasts, by as much as 2.70 log2 units. Cells on rigid substrate should therefore be more sensitive to inhibitors of cystine import or of GPX4 than the same cells on compliant substrate; the difference should be rescued by forcing SLC7A11 expression or by supplying cell-permeable glutathione precursors, and it should track lipid peroxidation rather than general cytotoxicity [42,43]. Because drug-tolerant persister states depend on the same axis [27], the test is worth doing in the persister setting as well. If it holds, matrix rigidity becomes a candidate marker for tumor regions likely to respond to agents that kill through oxidative damage; if it fails, the transcriptional change is buffered downstream, which would be equally worth knowing.

## Materials and methods

### Dataset identification and curation

The GEO DataSets database [28] was queried through the E-utilities API for transcriptome studies of human breast cancer cells cultured on substrates of defined stiffness. Candidate series were screened manually against four requirements: a human breast cancer line, at least two substrate conditions differing in modulus, at least three replicates per condition, and processed expression data deposited in a usable form. A sample-level design table recording platform, accession, species, cell line, substrate, modulus, replicate, inclusion status, and the reason for any exclusion is provided as S1 Table. Gene symbols were reconciled across platforms before any cross-dataset comparison, since the microarray series carry legacy symbols and the RNA-seq series current HGNC symbols for the same genes; CCN2 and CCN1 were mapped to CTGF and CYR61 respectively. The eleven series screened were GSE93529 [29], GSE107063 [30], GSE83366 [31], GSE182606 [32], GSE205816 [33], GSE203651 [44], GSE255827, GSE255829 and GSE255831 [45], and GSE274519 and GSE92653, the last two of which carry no linked publication and are cited by accession.

For GSE93529, the GEO record and the corresponding publication disagree on the moduli used; the GEO record (0.7 kPa hydrogel versus plastic) was taken as authoritative for the deposited samples, and this is noted in S1 Table.

### Quality control

Three diagnostics were computed per dataset before any differential testing. Minimum within-group Spearman correlation across all replicate pairs; condition separation, defined as the median within-group correlation minus the median between-group correlation; and, for RNA-seq datasets, the number of genes detected per sample at matched depth. Datasets were retained only where the minimum within-group correlation reached 0.9 and separation reached 0.02. Counts for RNA-seq datasets were summed across duplicate gene identifiers and converted to log2 counts per million before correlation. Principal component analysis was performed on log-transformed expression for each retained dataset.

### Differential expression

GSE83366 was processed from the deposited non-normalized Illumina file with the limma [46] function read.ilmn, background-corrected and quantile-normalized with neqc [47], and annotated with the illuminaHumanv4.db package rather than the GEO platform annotation. Because that package returns current gene symbols, CCN2 and CCN1 were mapped back to the aliases CTGF and CYR61 for display alongside the other contrasts. GSE93529 was analyzed from the submitter-deposited RMA values in the series matrix; local RMA could not be completed in this environment, and the deposited values were used instead. GSE107063 expression was taken from the submitter supplementary table accompanying the publication.

For all array datasets, linear models were fitted per probe with substrate condition as the sole factor and moderated t statistics computed with eBayes(trend = TRUE, robust = TRUE). P values were adjusted by the Benjamini-Hochberg procedure [48] within each contrast. Probes were collapsed to gene symbols by one of two rules, applied consistently within each analysis and stated here because they differ. For the module analyses and Figs 2 to 4, log2 fold changes were averaged across probes mapping to the same symbol and the smallest adjusted P value retained. For the quadrant analysis and Fig 5, the single probe with the smallest raw P value was retained per symbol. Both rules were applied to all contrasts alike within their respective analyses. Throughout, a positive log2 fold change denotes higher expression on the stiffer or more rigid substrate. Five contrasts were analyzed: MDA-MB-231 0.7 kPa to plastic; MDA-MB-453 5 to 30 kPa; MDA-MB-453 5 kPa to glass; and MDA-MB-231 and HeLa cell-derived matrix to plastic.

### Module definitions and testing

The NRF2 battery (14 genes), YAP/TEAD targets (9 genes) and ferroptosis effectors (6 genes) were specified from the literature before analysis and were not modified after results were seen. Module behavior was tested by comparing the moderated t statistics of module members against all remaining measured genes with a two-sided Mann-Whitney U test, separately in each contrast. Antioxidant response element dependence was assigned a priori to three tiers: pathway components (NFE2L2, KEAP1, CUL3, MAFG; 4 genes), moderately dependent genes (8 genes), and strongly dependent genes (11 genes). Tier means were compared across contrasts, and pathway components were compared with strongly dependent targets by Mann-Whitney U test on pooled log2 fold changes.

### Proliferation control

Two independent controls were applied. A proliferation pool was assembled from five MSigDB Hallmark sets [49] (E2F Targets, G2-M Checkpoint, Myc Targets V1 and V2, Mitotic Spindle), giving 671 unique genes, and module tests were repeated with these genes removed. Separately, each gene’s statistic was regressed on module membership and mean expression level by ordinary least squares, and module tests were repeated on the residuals, with a per-contrast proliferation coefficient recorded from the same model.

### Quadrant partition and over-representation analysis

Genes reaching FDR < 0.05 in both contrasts of a comparison were assigned to quadrants by the sign of their log2 fold change in each. Ribosomal protein, mitochondrial, and small nuclear or nucleolar RNA symbols were removed by regular expression before partitioning (159 symbols), leaving 16,953 genes measured in all three contrasts as the background for every enrichment test. Overlap between the two MDA-MB-453 arms was assessed within each quadrant by Jaccard index and a one-sided hypergeometric test against that background. Because the resulting gene sets are discrete rather than ranked, enrichment was tested by over-representation rather than pre-ranked gene set enrichment analysis [50]. Each set was tested by one-sided hypergeometric test against MSigDB Hallmark 2020 [49], GO Biological Process 2023 [51] and TRRUST Transcription Factors 2019 [52], restricted to terms with 5 to 600 background genes and at least 2 overlapping genes, with Benjamini-Hochberg correction applied within each library and quadrant. The proliferation-controlled analysis removed the 671 pool genes from both the query set and the background; the five Hallmark sets defining the pool were excluded from that comparison to avoid circularity.

### Human tumor cohorts

Expression for 1,097 TCGA-BRCA primary tumor aliquots was taken from the Xena HiSeqV2 matrix [34], and for 1,980 METABRIC tumors [36,37] from the Illumina HT-12 profile retrieved through the cBioPortal API [53,54]. Module scores were computed within each cohort as the mean z score across member genes. Three NRF2 scores were specified. The first used all fourteen battery genes tested in culture. The second used the ten measurable antioxidant response element-dependent members (SLC7A11, NQO1, TXNRD1, SRXN1, GCLM, GCLC, HMOX1, OSGIN1, G6PD and PGD), omitting TALDO1, GSR, ME1 and AKR1C1. The third, used as the primary outcome, is a seven-gene tumor-intrinsic score (SLC7A11, GCLM, NQO1, TXNRD1, SRXN1, OSGIN1, GCLC): the strongly dependent tier of Fig 4A with HMOX1 removed because it is substantially expressed by tumor-associated macrophages and AKR1C1 removed because it is unreliably detected. It was chosen on these grounds rather than on its performance. All three scores were carried through every test in both cohorts, and all results are tabulated in S1 Table; the text reports the primary score and notes where the three diverge. The matrix stiffness proxy was a thirteen-gene collagen and lysyl oxidase score (LOX, LOXL1, LOXL2, LOXL3, LOXL4, COL1A1, COL1A2, COL5A1, COL11A1, FN1, POSTN, TNC, PLOD2). It indexes desmoplastic content; it is not a measurement of tissue modulus, and it is not a validated stromal deconvolution instrument such as ESTIMATE [55]. Hypoxia was scored from ten canonical HIF1A targets (CA9, VEGFA, SLC2A1, LDHA, PGK1, ADM, NDRG1, P4HA1, BNIP3, ALDOA) rather than from a validated metagene such as that of Buffa and colleagues [35]. Epithelial content was scored from EPCAM, KRT7, KRT8, KRT18, KRT19 and CDH1, and myeloid content from PTPRC, CD68, CD163, AIF1, ITGAM, CSF1R, LYZ, MRC1 and MSR1.

Associations were measured by Spearman correlation and by ordinary least squares on z scored predictors, regressing the NRF2 score on the stiffness proxy, hypoxia, epithelial content and myeloid content. Partial correlations were obtained by rank transforming every variable, residualizing outcome and predictor on the covariates by least squares, and correlating the residuals. The covariate set was selected in TCGA after the unadjusted null had been observed; it was then fixed and applied unchanged to METABRIC, which served as the replication cohort with the stiffness coefficient as the pre-specified endpoint. Quartile comparisons contrasted the top and bottom quartiles of the proxy by Mann-Whitney U test, with Cohen’s d as the effect size. Subtype analyses used PAM50 calls and immunohistochemical receptor status. Extreme-group testing was exploratory: seventeen comparisons were run in TCGA across the three NRF2 scores and five strata (quartile, decile, high-epithelial quartile, high-epithelial decile, and PAM50 or receptor subgroup). None was corrected for multiplicity and all seventeen are reported in S1 Table. Only the pre-specified quartile contrast on the primary score is carried into the replication cohort and into the text.

Single-cell and spatial data came from the breast cancer atlas of Wu and colleagues [38]. For each tumor, counts were summed across cells within a compartment and then converted to counts per million and log transformed, giving one pseudobulk profile per tumor for cancer epithelial cells and one for cancer-associated fibroblasts. Tumors contributing fewer than 20 cells to a compartment were dropped, and module genes were required to be present in at least 70% of pseudobulk profiles. The NRF2 score was computed in malignant cells and the matrix score in fibroblasts, so the two derive from disjoint cell populations and cannot be coupled by tissue composition. For the six spatial sections, spot-level scores were computed in the same way, with module genes required in at least 10% of in-tissue spots, malignant-dominant spots defined as the top 40% of the epithelial score, and a neighborhood matrix score computed as the mean across spots within 1.5 spot spacings, excluding the spot itself. Spot-level models regressed the NRF2 score on the neighborhood matrix score with epithelial, myeloid and hypoxia scores, log total counts, genes detected and neighborhood depth as covariates and section as a fixed effect, with variables z scored within section and standard errors clustered by section. Ribosomal, mitochondrial and small nuclear or nucleolar RNA genes were excluded from all single-cell and spatial scoring.

## Software and availability

Analyses were performed in Python 3.11 with pandas [56], NumPy [57], SciPy [58], statsmodels [59] and matplotlib [60], and in R with limma [46], oligo and illuminaHumanv4.db. Gene set libraries were retrieved from Enrichr [61]. The sample design table, quality control metrics, differential expression for all five contrasts, gene lists, enrichment output, the proliferation-control comparison and the human cohort models are provided in S1 Table.

## Acknowledgments

### Data availability

All data analyzed in this study are publicly available. The expression series are in the NCBI Gene Expression Omnibus under accessions GSE93529, GSE107063, GSE83366, GSE182606, GSE205816, GSE203651, GSE255827, GSE255829, GSE255831, GSE274519 and GSE92653. The breast carcinoma cohort is available from the Genomic Data Commons as TCGA-BRCA, and the METABRIC cohort through cBioPortal as study brca_metabric. Single-cell data for the breast cancer atlas are in the Gene Expression Omnibus under GSE176078, and the matched spatial transcriptomic data in Zenodo under doi:10.5281/zenodo.4739739. The sample-level design table, differential expression for all five contrasts, gene lists, enrichment output, the proliferation-control comparison and the human cohort models and extreme-group tests are provided in S1 Table. Analysis code is deposited at [mine will be available soon] and archived at [Zenodo DOI: will be available soon].

## Supporting information

**S1 Fig. Enrichment of each quadrant across three gene set libraries.** Over-representation results for the four quadrants defined in Fig 5A, tested against MSigDB Hallmark 2020 (A), GO Biological Process 2023 (B) and TRRUST Transcription Factors 2019 (C), each against a background of 16,953 genes measured in all three contrasts. Each panel shows the two most significant terms per quadrant, to a maximum of eight, and carries its own color scale, because the libraries differ by an order of magnitude in the significance they reach. Dot area gives the number of overlapping genes and a black outline marks FDR < 0.05. Terms reaching FDR < 0.05 were, for Hallmark, 4 in Q1, 16 in Q2, 13 in Q3 and 6 in Q4; for GO biological process, 72, 59, 16 and 101; and for TRRUST, 2 in Q1, 26 in Q2 and none in Q3 or Q4. Leading Hallmark terms were E2F targets in Q1 (FDR 5.4 × 10□²¹), cholesterol homeostasis in Q2 (7.6 × 10□□), xenobiotic metabolism in Q3 (3.8 × 10□□) and MYC targets V2 in Q4 (1.8 × 10□²□). Leading GO terms were mRNA splicing via spliceosome in Q1 (2.8 × 10□□), cholesterol biosynthetic process in Q2 (3.1 × 10□□), tricarboxylic acid metabolic process in Q3 (8.8 × 10□³) and ribosome biogenesis in Q4 (1.7 × 10□²□).

**S2 Fig. Enrichment with proliferation genes removed.** The analysis of Fig 5G and S1 Fig repeated after removing 671 proliferation genes, pooled from five MSigDB Hallmark sets, from both the query sets and the background. The five Hallmark sets defining the pool were excluded from the comparison to avoid circularity. (A-D) The eight leading terms of each quadrant, with FDR before and after removal joined by a line; the dashed line marks FDR = 0.05. Proliferation genes made up 59 of 300 transcripts in Q1 (19.7%), 9 of 246 in Q2 (3.7%), 3 of 158 in Q3 (1.9%) and 44 of 247 in Q4 (17.8%). All eight Q3 terms held or strengthened. In Q1 the cell-cycle terms fell below significance, leaving only an mRNA splicing program (FDR = 1.1 × 10□□). Across 32 term comparisons, 28 held, 3 weakened, and 1 was lost.

**S3 Fig. Diagnostics for the human cohort analysis.** (A) Spearman correlation of the matrix stiffness proxy with epithelial, myeloid and hypoxia scores in TCGA and METABRIC. The proxy tracks myeloid content and hypoxia but not epithelial content, which is the structure that suppresses the association in Fig 6A. (B) Log2 fold change in culture across the five contrasts, for the eight battery genes detectable in single-cell data against the six that are not (means −0.38 and −0.76; Mann-Whitney P = 0.024). The effect concentrates in the genes that sparse assays cannot measure. (C) Spatial estimate in 6,241 malignant-dominant spots across six sections as sequencing-depth covariates are added; the unadjusted positive association does not survive control for spot depth, genes detected and neighborhood depth.

**S1 Table. Design table, quality control, differential expression, enrichment and human cohort results.** Single workbook of sixteen sheets. The sample-level design table (S1) records platform, accession, title, species, cell line, substrate, modulus, replicate, inclusion status and the reason for every exclusion, for all 137 samples deposited across the eleven series screened, and notes the modulus discrepancy in the GSE93529 record. Quality control occupies four sheets: the summary metrics behind Fig 1A and 1B (S2), every within-group and between-group replicate correlation (S3), the principal component coordinates behind Fig 1D to 1F for all four retained arms (S4), and the per-sample gene detection counts behind Fig 1C (S5). S6 lists the data sources and filters, and S7 gives differential expression for all five contrasts across 23,924 genes, flagging the 16,953 that constitute the over-representation background. S8 to S11 give the quadrant gene lists, the overlap statistics between the two MDA-MB-453 arms, the full over-representation output for every quadrant and library, and the term-by-term comparison before and after proliferation control. S12 reports the multivariable coefficients and partial correlations for both cohorts on all three NRF2 scores, S13 the seventeen extreme-group comparisons across those scores and strata, S14 the single-cell compartment scores, and S15 and S16 the spatial depth cascade and per-section statistics.

