## Supplementary Fig's for "Substrate rigidity suppresses the NRF2 antioxidant program in breast cancer cells"

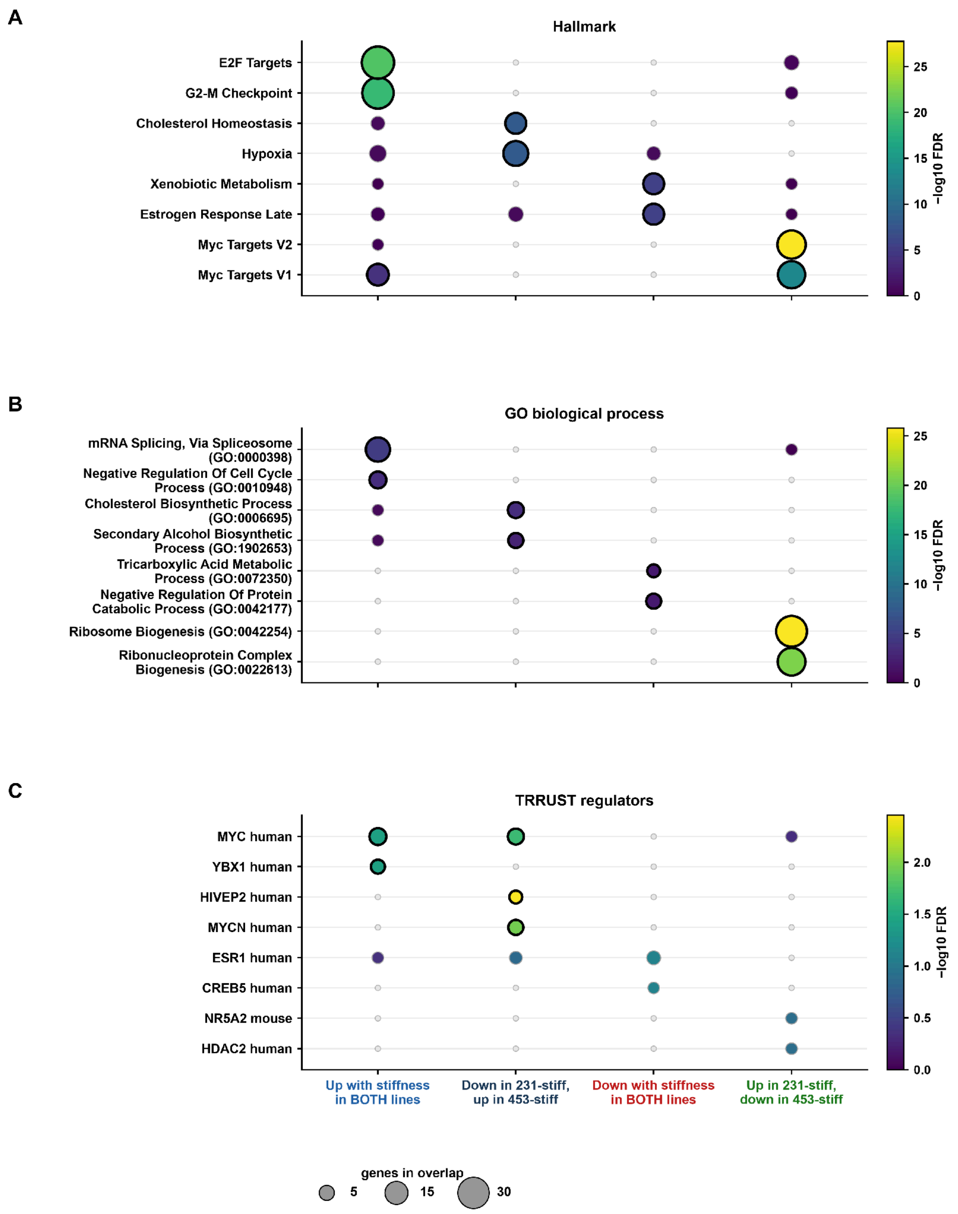


**S1 Fig. Enrichment of each quadrant across three gene set libraries.** Over-representation results for the four quadrants defined in Fig 5A, tested against MSigDB Hallmark 2020 (A), GO Biological Process 2023 (B) and TRRUST Transcription Factors 2019 (C), each against a background of 16,953 genes measured in all three contrasts. Each panel shows the two most significant terms per quadrant, to a maximum of eight, and carries its own color scale, because the libraries differ by an order of magnitude in the significance they reach. Dot area gives the number of overlapping genes and a black outline marks FDR < 0.05. Terms reaching FDR < 0.05 were, for Hallmark, 4 in Q1, 16 in Q2, 13 in Q3 and 6 in Q4; for GO biological process, 72, 59, 16 and 101; and for TRRUST, 2 in Q1, 26 in Q2 and none in Q3 or Q4. Leading Hallmark terms were E2F targets in Q1 (FDR $5.4 \times 10^{-21}$), cholesterol homeostasis in Q2 ($7.6 \times 10^{-9}$), xenobiotic metabolism in Q3 ($3.8 \times 10^{-6}$) and MYC targets V2 in Q4 ($1.8 \times 10^{-28}$). Leading GO terms were mRNA splicing via spliceosome in Q1 ($2.8 \times 10^{-5}$), cholesterol biosynthetic process in Q2 ($3.1 \times 10^{-4}$), tricarboxylic acid metabolic process in Q3 ($8.8 \times 10^{-3}$) and ribosome biogenesis in Q4 ($1.7 \times 10^{-26}$).

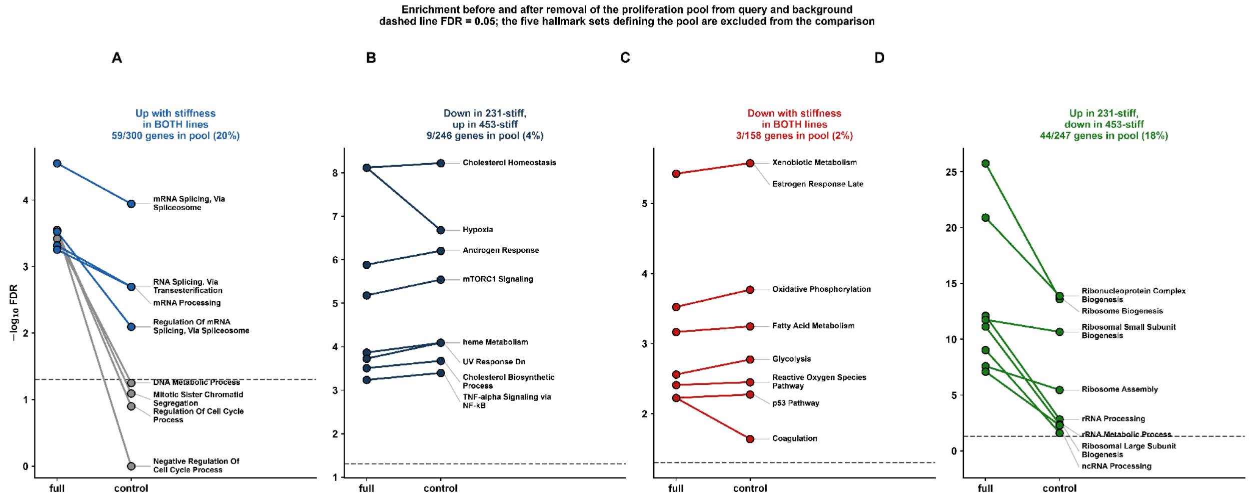


**S2 Fig. Enrichment with proliferation genes removed.** The analysis of Fig 5G and S1 Fig repeated after removing 671 proliferation genes, pooled from five MSigDB Hallmark sets, from both the query sets and the background. The five Hallmark sets defining the pool were excluded from the comparison to avoid circularity. (A-D) The eight leading terms of each quadrant, with FDR before and after removal joined by a line; the dashed line marks FDR = 0.05. Proliferation genes made up 59 of 300 transcripts in Q1 (19.7%), 9 of 246 in Q2 (3.7%), 3 of 158 in Q3 (1.9%) and 44 of 247 in Q4 (17.8%). All eight Q3 terms held or strengthened. In Q1 the cell-cycle terms fell below significance, leaving only an mRNA splicing program (FDR = $1.1 \times 10^{-4}$). Across 32 term comparisons, 28 held, 3 weakened and 1 was lost.

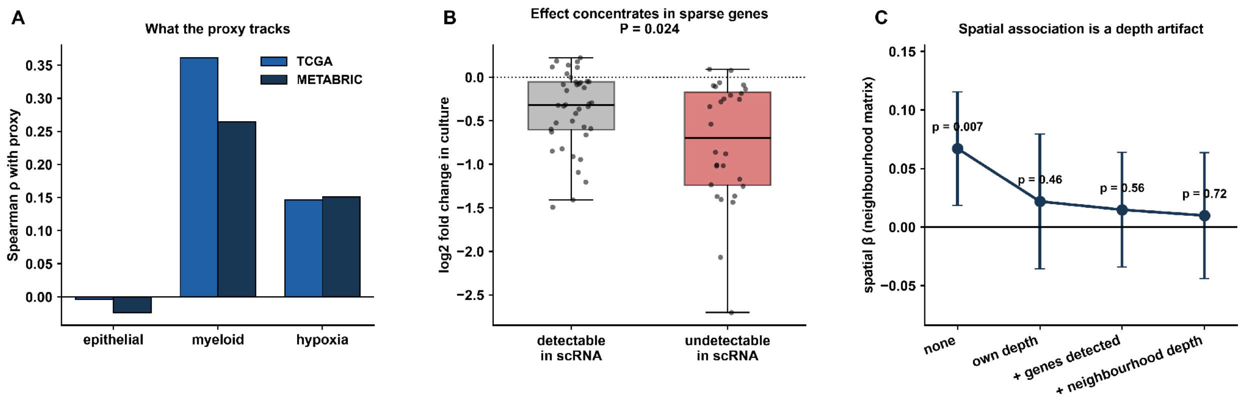


**S3 Fig. Diagnostics for the human cohort analysis.** (A) Spearman correlation of the matrix stiffness proxy with epithelial, myeloid and hypoxia scores in TCGA and METABRIC. The proxy tracks myeloid content and hypoxia but not epithelial content, which is the structure that suppresses the association in Fig 6A. (B) Log2 fold change in culture across the five contrasts, for the eight battery genes detectable in single-cell data against the six that are not (means −0.38 and −0.76; Mann-Whitney P = 0.024). The effect concentrates in the genes that sparse assays cannot measure. (C) Spatial estimate in 6,241 malignant-dominant spots across six sections as sequencing-depth covariates are added; the unadjusted positive association does not survive control for spot depth, genes detected and neighborhood depth.

**S1 Table. Design table, quality control, differential expression, enrichment and human cohort results.** Single workbook of sixteen sheets. The sample-level design table (S1) records platform, accession, title, species, cell line, substrate, modulus, replicate, inclusion status and the reason for every exclusion, for all 137 samples deposited across the eleven series screened, and notes the modulus discrepancy in the GSE93529 record. Quality control occupies four sheets: the summary metrics behind Fig 1A and 1B (S2), every within-group and between-group replicate correlation (S3), the principal component coordinates behind Fig 1D to 1F for all four retained arms (S4), and the per-sample gene detection counts behind Fig 1C (S5). S6 lists the data sources and filters, and S7 gives differential expression for all five contrasts across 23,924 genes, flagging the 16,953 that constitute the over-representation background. S8 to S11 give the quadrant gene lists, the overlap statistics between the two MDA-MB-453 arms, the full over-representation output for every quadrant and library, and the term-by-term comparison before and after proliferation control. S12 reports the multivariable coefficients and partial correlations for both cohorts on all three NRF2 scores, S13 the seventeen extreme-group comparisons across those scores and strata, S14 the single-cell compartment scores, and S15 and S16 the spatial depth cascade and per-section statistics. Supplied as a separate Microsoft Excel workbook (S1_Table.xlsx).
